# Hypobaric hypoxia drives citrate cycle reprogramming to suppress tumor progression

**DOI:** 10.1101/2025.08.15.669572

**Authors:** Yixing Gao, Xiuju He, E Guoji, Yan Wang, Shirui Huang, Dan Cai, Yumei Deng, Ping Gan, Rui Feng, Qian Liang, Ziyi Tang, Gang Xu, Jian Chen, Hongyu Xue, Qiutong Wang, Huanhuan Liu, Guibo Li, Shuang Zeng, Min Xie, Yuqi Gao, Jiao Gong, Erlong Zhang

## Abstract

High altitude residents present lower tumor incidence and mortality than plain residents, suggesting that hypobaric hypoxia displays protective effect against tumor development. The results of our studies showed that hypobaric hypoxia exposure significantly suppressed tumor progression in subcutaneous bearing and lung metastasis tumor models. With single-cell transcriptome analysis, we identified HIF-1α pathway was significantly downregulated in tumor tissues with high altitude hypoxia, leading to decreased glycolysis and angiogenesis and improved antitumor immune response. Mechanistically, systemic hypoxia rewired citric acid cycle, with enhanced CS and IDH2 expression and reduced OGDH and SUCLG2 expression, to induce α-ketoglutarate accumulation and succinate decline in tumor microenvironment, further mediating HIF-1α pathway inhibition. Moreover, hypobaric hypoxia treatment significantly improved the antitumor effects of adjuvant therapies (including chemotherapy, anti-angiogenic therapy and immunotherapy). Our findings reveal the role and mechanism of hypobaric hypoxia for tumor regression and yield new insights into tumor therapy.

## 1 Introduction

Tumor hypoxia was first discovered in 1955 by Thomlinson et al. from lung cancer tissues and has been confirmed widespread existence in a variety of solid tumors [1]. The appearance of hypoxic region in human solid tumors is due to rapid proliferation of tumor cells and the relative deficiency of blood distribution in the tumor mass, resulting in low oxygen levels in tumor cells, so-called hypoxic cells [2–3]. And decreased availability of oxygen in the tumor promotes therapeutic resistance and favors tumor progression and metastasis. Moreover, intra-tumor hypoxia is linked to decreased disease-free survival outcomes [4]. Furthermore, hypoxia regulate tumor malignant behaviors, including angiogenesis, tumor cell aggressiveness, metastasis, local recurrence and anti-tumor treatment outcomes [2–3].

However, different from the effect of local hypoxia in tumor tissues, systemic hypoxia, such as hypobaric hypoxia in high altitude, showed a beneficial role for tumor development. Many ecological studies reported that humans living at high altitude showed low incidence and reduced cancer mortality over a broad spectrum of cancer types, including lymphoma, breast cancer, cancer of the tongue, mouth, larynx, lung, and et al [5–6]. Oxygen-independent and oxygen-dependent mechanisms that might contribute to reduced cancer mortality at high altitude. Moreover, tumor metabolism and immune environment could be involved in this process. However, high altitude could increase the risk of some other tumors, such as carotid body tumors and skin cancer [5–6]. Recently, Arenillas et al. found that there was a broad convergence in genetic adaptation to hypoxia between natural populations and tumors [7], supporting high altitude adaptation could promote tumor progression. Therefore, there is a controversy about the effect of high altitude on tumor development.

With regard to distinct effects of hypoxia on tumor progression, studies investigating the role of oxygen supply on tumors has been performed. Hatfield et al. found that respiratory hyperoxia improved immune response in tumor environment to inhibit tumor progression by reversing hypoxia-adenosinergic immunosuppression [8]. More works supported that hyperbaric oxygen could promote cancer treatment by normalization of mechanics and blood vessels, suppressing cancer stem cell and metastasis, immunosuppression disruption and increased oxidative stress [9]. Additionally, the antitumor ability of adjuvant therapies could also be enhanced by hyperbaric oxygen [10–12]. Nevertheless, another work reported that normobaric hypoxia showed an opposite effect on lung cancer and colon cancer [13].

Recently, a study predicts the therapeutic role of hypobaric hypoxia environment for pan-cancer [14], which remains to be further determined. In this work, we evaluated the effect of hypobaric hypoxia on subcutaneous bearing and lung metastasis tumor models. Based on the results, the single cell transcriptome analysis from tumor tissues presented the characters of cell subtypes and gene expression across tumor environment. Our study testifies antitumor influence of hypobaric hypoxia and unveils that high altitude treatment leads to citrate cycle reprogramming in cancer cells to promote HIF-1α downregulation. This discovery could not only bring new strategies for cancer treatment, but also open an avenue to interpret the low tumor risk in high altitude populations.

## 2 Materials and methods

### 2.1 Establishment and treatment for mice tumor models

The animal experimental protocols were approved by the Animal Care and Use Committee Guidelines of the Army Medical University (AMUWEC20245324). Special pathogen-free (SPF) C57 BL/6J mice weighing 18-20 g (aged 6 weeks) were purchased from Beijing Vitalriver Experimental Animal Technology Co., Ltd. For the subcutaneous syngeneic tumor model, Lewis Lung Carcinoma (LLC) cells (3.5 × 10^6^), colorectal cancer MC38 cells (1.0 × 10^6^) or melanoma B16 cells (2.0 × 10^6^) were injected subcutaneously into the right hind flanks of mice. Seven days later, the mice were randomized into groups fed at 300 m altitude above sea level (local altitude) or indicated altitude in the hypobaric hypoxia chamber. The implanted tumors were measured every two or three days, monitored by a digital caliper, and quantified by the modified ellipsoidal formula (tumor volumelJ=lJ1/2 (lengthlJ×lJwidth^2^)). Mice were killed and tumor tissues were harvested on Day 17 for LLC, Day 20 for MC38 and Day 19 for B16. For another syngeneic mouse tumor model used for pulmonary metastasis colonization assay, LLC cells (2.0 × 10^6^), MCA205 fibrosarcoma cells (1.0 × 10^6^) or B16 cells (0.3 × 10^6^) were injected through the lateral tail vein of C57 BL/6J mice. Mice were killed and lung tissues were harvested on Day 44 for LLC, Day 22 for MCA205 and Day 21 for B16.

For HIF-1α related treatment, the mice were treated with saline control, 100 mg/Kg DMOG (MCE, Cat# HY-15893), or 60 mg/Kg PX-478 (MCE, Cat# HY-10231) three times a week by intraperitoneal injection. Mice were killed and tumor tissues were harvested on Day 25. For the TCA cycle metabolite treatment, the mice were treated with saline control, 600 mg/Kg dimethyl 2-oxoglutarate (DM-KG) (MCE, Cat# HY-44134), or 100 mg/Kg dimethyl succinate (DM-S) (MCE, Cat# HY-Y0808) on workdays by intraperitoneal injection. Mice were killed and tumor tissues were harvested on Day 19. For drugs combined therapies, the mice were treated with saline control, 5mg/Kg bevacizumab (Bev) (Ankeda), 1.5 mg/Kg CDDP (Ailikang), or 10 mg/Kg anti PD-L1 (BioXCell, Cat# BE0101) once two days. Mice were killed and tumor tissues were harvested on Day 23.

### 2.2 Preparation of cell suspensions from mouse tumor tissues

To prepare single-cell suspensions, three subcutaneous LLC tumor tissues from normobaric normoxia or hypobaric hypoxia were cut into pieces and incubated in the 15mL-tube with enzyme mix containing 250 U/mL collagenase I (Sangon Biotech, Cat# A004194-0001), 100 U/mL collagenase IV (Sangon Biotech, Cat# A004186-0001), and 10 U/mL DNase I (Sigma-Aldrich, Cat# D7291-2MG) in RPMI 1640 medium at 37 lJ for 15 minutes, accompanied by manual blowing every 5 minutes. The homogenate was filtered through a 70 μm strainer and supplemented with fetal bovine serum to terminate digestion. The residual particles were incubated with another enzyme mix containing 200 U/mL collagenase I, 400 U/mL collagenase II (Sangon Biotech, Cat# A004174-0001), and 10 U/mL DNase I in RPMI 1640 medium at 37 lJ for 15 minutes and then filtered through a 70 μm strainer, accompanied by manual blowing every 5 minutes. Afterward, all the cell suspension was centrifuged at 500×g for 10 minutes at 4 lJ, added with 4 mL of erythrocyte lysis buffer, and incubated on ice for 5 minutes for erythrocyte lysis. After washing with PBS containing 1.5% FBS and 20 mM EDTA, the cells were resuspended in 1×PBS (calcium and magnesium free) containing 0.04% weight/volume BSA (400 µg/mL).

### 2.3 single cell RNA-sequencing (scRNA-seq)

Tumor sample libraries were constructed following the manufacturer’s protocol for the DNBelab C Series High-throughput Single-cell RNA Library Preparation Set V3.0 (MGI, https://en.mgi-tech.com/products/solution/3/). Briefly, cells undergo lysis within microfluidic droplets, where released mRNA molecules become immobilized and uniquely labeled with barcodes. After reverse transcription, complementary DNA (cDNA) synthesis occurs, followed by amplification. These amplified cDNA products are then fragmented to generate inserts ranging from 400 to 600 base pairs (bp). Library construction for cDNA and oligo components featured distinct read structures: cDNA libraries incorporated a 41-bp Read 1 (containing a 20-bp cell barcode and a 10-bp unique molecular identifier, UMI), a 100-bp Read 2, and a 10-bp sample index. Oligo libraries were constructed with a 26-bp Read 1, a 42-bp Read 2, and a 10-bp sample index.

### 2.4 scRNA-seq data analysis

#### Processing and Annotation of scRNA-seq Data

Raw scRNA-seq data were processed automatically using the BGI analytical pipeline. Sequencing reads underwent sequential computational steps, including barcode demultiplexing, adapter sequence filtering, reference genome alignment, PCR duplicate removal, and gene expression quantification. The DoubletFinder algorithm was employed to remove doublets [15]. To ensure rigorous quality control, cells exhibiting fewer than 500 unique molecular identifiers (UMIs) and genes detected in fewer than three cells were systematically excluded. Subsequent data processing using Seurat’s standard parameters [16]. Comprehensive details of the sequencing methodologies and analytical procedures are available in Table S1.

#### Differential Gene Expression and GO Enrichment Analysis

Following normalization, the six samples were seamlessly integrated using Seurat’s ‘merge’ function, and batch effects were effectively mitigated with Harmony [17]. Differential expression analysis was executed via the ‘FindMarkers’ function in Seurat. Genes exhibiting an average log-fold change exceeding 0.25 and an adjusted p-value under 0.05 were classified as differentially expressed. The resulting DEG lists served as direct input for the ‘clusterProfiler’ R package [18–19] to conduct Gene Ontology (GO) enrichment analysis. Benjamini-Hochberg correction controlled for multiple testing, and GO terms with an adjusted *p*-value < 0.05 were deemed statistically significant. Enrichment outcomes were clearly visualized.

#### GSVA annalysis

GSVA analysis was performed to quantify pathway activity scores across cell clusters, using the Hallmark gene sets from MSigDB (https://www.gsea-msigdb.org/gsea/msigdb/index.jsp) with default parameters [20]. Pathway scores were then correlated with communication strength to identify signaling mechanisms driving inter-sample differences, and results were visualized via heatmaps highlighting enriched pathways under hypobaric hypoxia.

#### Reguron Annalysis

pySCENIC [21–22] analysis was conducted to infer gene regulatory networks and identify key transcription factors (TFs) across cell types, using the standard workflow with default parameters. The results revealed cell-type specific regulatory programs, highlighting enriched TFs and their target genes, which were visualized via heatmaps and network diagrams to elucidate mechanisms driving sample heterogeneity.

### 2.5 Determination of metabolite and reactive oxygen species (ROS) content

For metabolite measurement from tumor tissues, 70-90 mg of frozen/fresh tissue was homogenized in BeyoLysis™ Buffer A for Metabolic Assay by Servicebio 24 homogenizer, and the supernatant was collected after centrifugation at 12,000g for 5 minutes at 4 lJ. The concentration of α-ketoglutarate (α-KG) and fumarate in the solution was then determined by the Amplex Red α-KG Assay Kit (Beyotime, Cat# S0323S) and Fumarate Assay Kit with WST-8 (Beyotime, Cat# S0517S) separately according to the manufacturer’s protocol for colorimetric assay without the deproteinization step. For succinate measurement, Succinic Acid Colorimetric Assay Kit (Elabscience, Cat# E-BC-K902-M) was used for colorimetric assay with the deproteinization step by Amicon® Ultra (Millipore, Cat# UFC8010). The ROS levels in tumor tissues were determined by a 96-well plate spectrophotometer (Multiskan GO, Thermo Scientific) using a reactive oxygen species assay kit (Bioswamp, BTK023). The whole process was performed according to the manufacturer’s instructions.

### 2.6 Cell culture

LLC cells, MC38 cells and B16 cells purchased from Zhongqiaoxinzhou Biotechnology and MCA205 cells purchased from TechiSun Biotechnology, and cultured with DMEM (LLC and MC38) or RPMI-1640 (B16 and MCA205) media supplemented with 10% fetal bovine serum (Hyclone, Cat# SH30406.05) and 1% penicillin/streptomycin at 37℃ in a 5% CO_2_ humidified incubator.

### 2.7 Cellular transfection experiments

Each group of cells was seeded and cultured overnight before transfection. For the transient overexpression of target genes, overexpression plasmids or control plasmids (GenePharma) were transfected into the indicated cells using the Lipofectamine 3000 transfection reagent (Invitrogen). For the knockdown of target genes, the specific siRNA or scrambled siRNA (Scr) (GenePharma) were transfected into the indicated cells using the Lipofectamine RNAiMAX transfection reagent (Invitrogen).

### 2.8 RNA isolation and RT-qPCR

The total RNA from cells or tumor tissue was extracted with Trizol and chloroform, and purified with isopropanol and ethanol. Subsequently, reverse transcription was performed using a PrimeScript™ RT reagent Kit with gDNA Eraser (TaKaRa). Finally, the real-time polymerase chain reaction (qPCR) was run in a CFX96 Real-time System (Bio-Rad). The sequences of the primers for qPCR are listed in Supplementary Table 2.

### 2.9 Protein lysate preparation and western blotting

The total protein was extracted by RIPA lysis buffer (Beyotime) with Protease Inhibitors and Phosphatase Inhibitor Cocktails (MCE). Protein lysates were quantified with BCA assay, normalized, denatured (95 °C), and resolved on a 10% Tris-Glycine SDS-PAGE followed by immunoblotting. The antibodies used in the study are listed in Supplementary Table 3.

### 2.10 Hematoxylin and eosin (H&E) staining

Paraffin-embedded tissues were sectioned (4 μm), dewaxed using xylene, and rehydrated through a serial alcohol gradient. After washing with 1lJ×lJPBS, the slides were stained with hematoxylin and eosin (Beyotime) and dehydrated through increasing concentrations of ethanol and xylene.

### 2.11 Immunohistochemistry

The samples followed the steps of deparaffinization in xylene, rehydration through an ethanol series, antigen retrieval (10 mM sodium citrate buffer, pH 6.0), treatment with 3% hydrogen peroxide to inhibit endogenous peroxide activities, and nonspecific binding was blocked with normal goat serum. Tissue slides were then incubated with primary antibodies. For IHC staining, tissue sections were incubated with the Biotin-labeled Goat Anti-Rabbit IgG (Beyotime, Cat# A0279) and the Streptavidin, Peroxidase, R.T.U. (Vector, Cat# SA-5704-100), followed by detection using the ImmPACT® DAB Substrate Kit, Peroxidase (Vector, Cat# SK-4105) and counterstaining with hematoxylin. The antibodies used in the staining are listed in Supplementary Table 3.

### 2.12 Luciferase promoter assay

A dual luciferase reporter assay system (Promega) was used according to the manufacturer’s instructions. Briefly, cells were plated into 24-well plates and incubated at 37 °C in a 5% CO_2_ incubator overnight. Subsequently, a mixture containing Lipofectamine 3000 transfection reagent (Invitrogen), luciferase *Idh2* or *Ogdh* promoter (∼2000 bp upstream of the start site) vector (GenePharma), Renilla vector, and *Ybx1* vectors or *Tcf12* specific siRNA and scramble siRNA was added. Luciferase and Renilla signals were measured 48 h after transfection.

### 2.13 Statistical analysis

Statistical analysis was carried out using R (version 3.6.0), SPSS Statistics (version 29.0) and GraphPad Prism (version 9.0). The experiments were analyzed using independent t-tests for two groups or one-way ANOVA for multiple groups. Tumor volume from repeated-measure designs were analyzed using two-way ANOVA (or mixed model). Data were presented as the mean ± SEM. *p*<0.05 was regarded as significantly different. All experiments were independently carried out at least three times.

## 3 Results

### 3.1 Hypobaric hypoxia suppressed tumor progression

Mice subcutaneous bearing and lung metastasis tumor models were performed to evaluate the effect of hypobaric hypoxia on tumor progression. The LLC cells, MC38 cells and B16 cells were implanted into subcutaneous tissues and the mice were killed at corresponding dates (Figure 1A) and the tumor tissues were collected and imaged (Figure 1B). The growth curve from subcutaneous tumors showed that hypobaric hypoxia significantly inhibited the progression of these three tumor models in vivo. Moreover, the volume and weight of hypobaric hypoxia treated mice remarkably decreased compared with that of mice under normobaric normoxia (Figure 1B). As to LLC cells, MCA205 and B16 cells lung metastasis models, the mice were killed and the lung tissues were collected, imaged and weight (Figure S1A). Tumor regression was observed in mice with established lung tumors treated with hypobaric hypoxia and tumor nodules reduced noticeably (Figure 1C).

**Figure 1.**
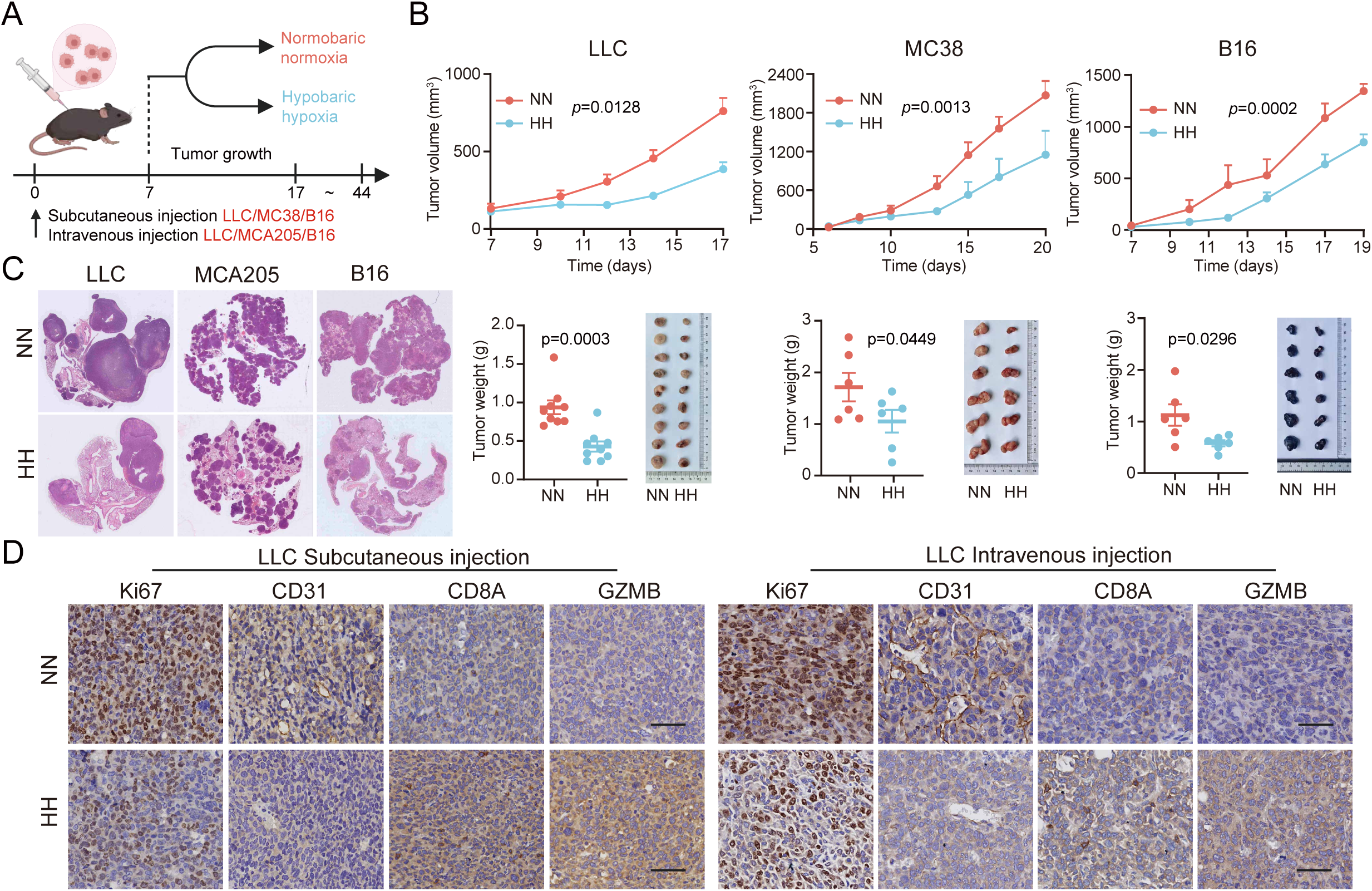
Hypobaric hypoxia exposure suppressed tumor progression by impairing malignant phenotypes. (A) An overview of the workflow for the establishment and following treatment about mice tumor models. (B) The growth curve, weight and images of tumor tissues from LLC, MC38 and B16 subcutaneous tumor models. (C) The transverse sections of lung tissues from LLC, MCA205 and B16 metastatic tumor models. (D) Immunohistochemical staining of Ki67, CD31, CD8A and GZMB from LLC subcutaneous and intravenous tumor models. Data represent mean ± SEM. Scale bar=50 μm.

To clarify antitumor effects from high altitude exposure, the proliferation ability, vascular distribution and immune related proteins were observed in the slices from tumor tissues. The results showed a significant decrease in the expression of Ki67 and CD31, suggesting inhibition of cell proliferation and vascular formation. Moreover, the expression of CD8A and GZMB significantly increased in tumor tissues under hypobaric hypoxia, indicating tumor immune environment was improved (Figure 1D and S1B-C).

### 3.2 Hypobaric hypoxia promoted tumor suppressive effect through downregulation of HIF-1**α** pathway

To elucidate the mechanisms underlying tumor suppression under hypobaric hypoxia, we performed single-cell transcriptome analysis for the subcutaneous LLC tumor-bearing mice model under plain and 5,800m plateau, incorporating three biological replicates per group. Following stringent quality control and filtering, our analysis yielded a total of 62,127 high-quality cells. These cells resolved into 16 distinct clusters, six robust tumor cell populations characterized by *Krt8* expression, seven diverse immune cell populations identified by *Ptprc* expression (including three macrophage subsets and one dendritic cell subset marked by *Itgam* expression), two endothelial cell populations defined by *Pecam1* expression, and a single fibroblast population (Figure 2A-D, Figure S2A). Pronounced upregulation was observed in genes associated with translation, while notable downregulation occurred in genes linked to immune cell differentiation (Figure S2B). Among these clusters, the proportion of myeloid-derived suppressor cells in tumor tissues was decreased by hypobaric hypoxia treatment (Figure S2E-G).

**Figure 2.**
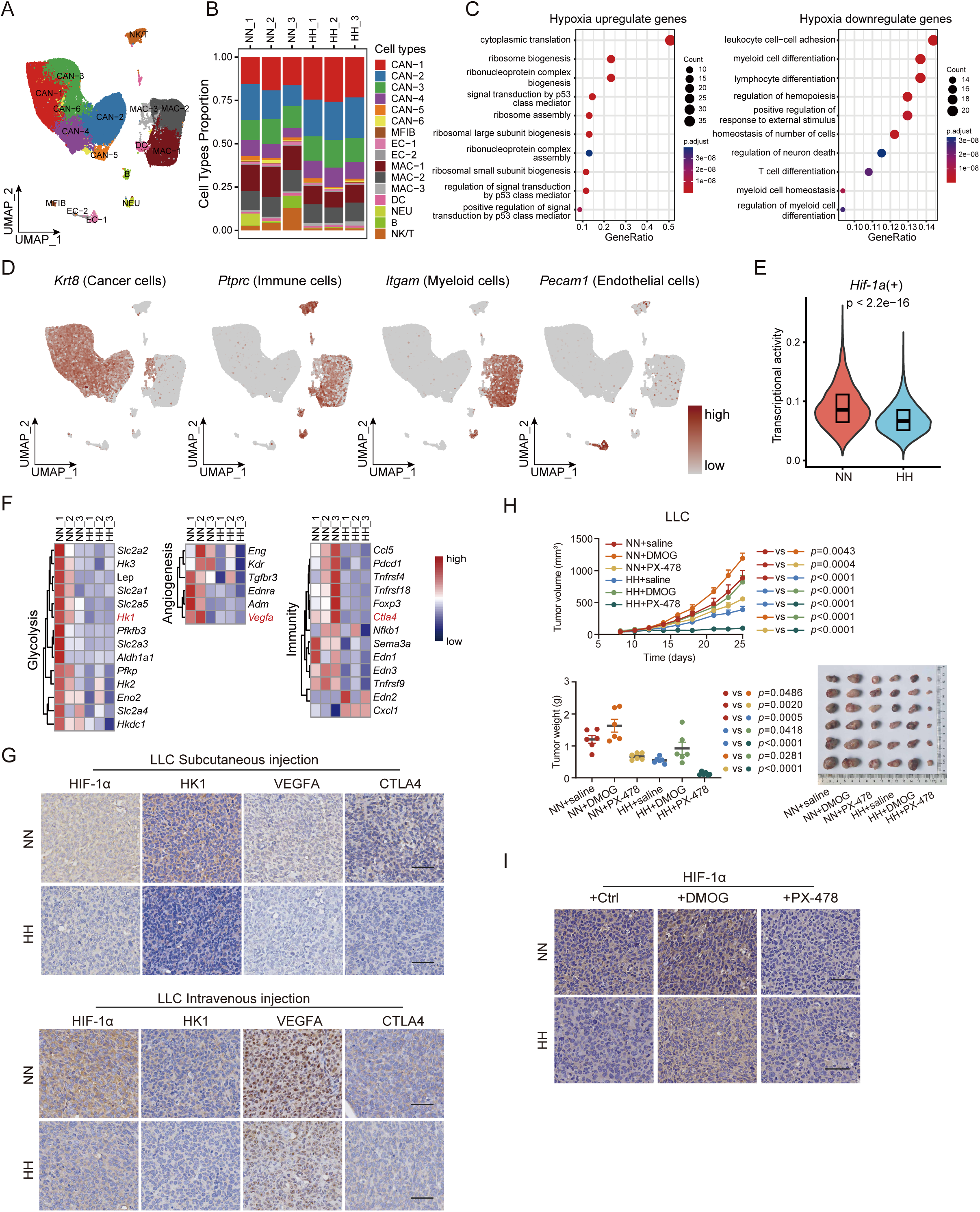
HIF-1α pathway was downregulated by hypobaric hypoxia in tumor environment. (A) The UMAP visualization of single-cell transcriptomes from subcutaneous LLC tumor tissues. (B) Proportional distribution of identified cell types. (C) Gene ontology biological process enrichment analysis of hypoxia-upregulated and hypoxia-downregulated genes. (D) Feature plot visualization of marker genes: *Krt8* (cancer cells), *Ptprc* (immune cells), *Itgam* (myeloid cells), *Pecam1* (endothelial cells). (E) Transcriptional activity of *Hif-1*α across all cells. (F) Expression profiles of key components within HIF-1α downstream genes about glycolysis, angiogenesis and immune response. (G) Immunohistochemical staining of HIF-1α, HK1, VEGFA and CTLA4 from LLC subcutaneous and intravenous tumor models. (H) The growth curve, weight and images of tumor tissues from LLC subcutaneous tumor models receiving saline, DMOG and PX478 treatment under plain and plateau. (I) Immunohistochemical staining of HIF-1α from LLC subcutaneous tumor models receiving saline, DMOG and PX478 treatment under plain and plateau. Data represent mean ± SEM. Scale bar=50 μm.

Our scRNA-seq analysis uncovered distinct context-dependent HIF suppression in tumor tissues bearing hypobaric hypoxia. Notably, context-specific HIF suppression was correlated tightly with reduced transcriptional activity (Figure 2E), rather than diminished *Hif-1*α expression itself (Figure S2C). And gene set variation analysis (GSVA) scores for hallmark gene sets presented a stark attenuation of hypoxia, angiogenesis and metabolism (S3A). This inhibition encompassed pivotal HIF-regulated processes, including impaired expression of glycolytic enzymes, diminished expression of vascular growth factors, improved immune effector functions (Figure 2F) and partially reduced expression with proliferation, invasion/metastasis and cancer stem cells (Figure S2D). Moreover, AddModuleScore analysis showed impaired hypoxia process (Figure S3B), which proved the phenomenon. Additionally, the expression of CA9, a marker for hypoxia evaluation, displayed no distinct change between plain and high altitude treatment, indicating that systemic hypoxia did not reduce oxygen supply within tumor environment and lead to the increased expression of HIF-1α (Figure S5B).

Based on the results of malignant phenotype and scRNA-seq analysis, we concluded that hypobaric hypoxia could inhibit glycolysis and vascular formation and improve tumor immunosuppressive microenvironment through repressing HIF-1α pathway. To validate this hypothesis, the expression of HIF-1α downstream proteins, differentially in scRNA-seq, was evaluated in the tumor tissues above. Along with the reduction of HIF-1α protein, the expression of glycolytic enzyme HK1, angiogenesis related protein VEGFA, and immune checkpoint protein CTLA4, was significantly decreased in the tumor tissues with high altitude exposure (Figure 2G, S2C and S4A-B).

Furthermore, the LLC subcutaneous bearing tumors were treated by dimethyloxaloyl glycine (DMOG), a HIF-1α agonist, and PX478, a HIF-1α inhibitor. And DMOG could advance tumor progression under normoxia and hypoxia, but PX478 showed an opposite effect (Figure 2H). These results proved the oncogenic role of HIF-1α pathway and implied the suppressive effect of hypobaric hypoxia on tumor development was dependent on HIF-1α compromise. Moreover, the effect of hypobaric hypoxia on the expression of HIF-1α (Figure 2I) and Ki67, CD31 and CD8A (Figure S4C) was also revered by DMOG and enhanced by PX478.

### 3.3 Hypobaric hypoxia reprogrammed citrate cycle to impair HIF-1**α** pathway

To reveal the reason for HIF-1α reduction in tumor tissues under hypobaric hypoxia, we investigated the factors related to HIF-1α regulation [23]. The mRNA levels of *Hif-1*α and *Hif-2*α presented no alterations between normobaric normoxia and hypobaric hypoxia (Figure S5A), indicating that hypobaric hypoxia could influence the stabilization of HIF-1α protein instead of transcriptional regulation. However, the expression of VHL, FIH-1, PHD2 and p300 (Figure S5B) and the ROS content (Figure S5C) also displayed no changes upon high altitude exposure.

Interestingly, we observed dramatically reprogrammed expression for TCA related genes (Figure 3A) and markedly elevated scores (Figure 3B) for the TCA cycle pathway derived from scRNA-seq analysis, suggesting profound alterations in the TCA cycle within tumors under hypobaric hypoxia. And upregulated *Cs* and *Idh2*, which metabolized acetyl-CoA to α-KG, and downregulated *Ogdh* and *Suclg2*, which metabolized α-KG to succinate, was emerged in tumor cells (Figure 3C). With validation by immunohistochemical detection in tumor tissues, the expression of CS and IDH2 was increased, and the expression of OGDH and SUCLG2 was decreased under hypobaric hypoxia (Figure 3D, S5D-E). In addition, the upregulation of IDH3A and SDHA, was also involved the reprogrammed changes of TCA cycle metabolites (Figure S6A). These enzymatic shifts ultimately led to the accumulation of α-KG and a marked decrease of succinate (Figure 3E). Coincidently, the increased α-KG and decreased succinate exactly regulated the suppression of HIF-1α pathway and tumor progression under hypobaric hypoxia. Additionally, differentially from LLC tumors, the MC38 tumors showed decreased fumarate with upregulated SDHA and FH1 and the B16 tumors showed no succinate alteration with no obvious change of SDHA and FH1 under plateau exposure (Figure 3E and S6B).

**Figure 3.**
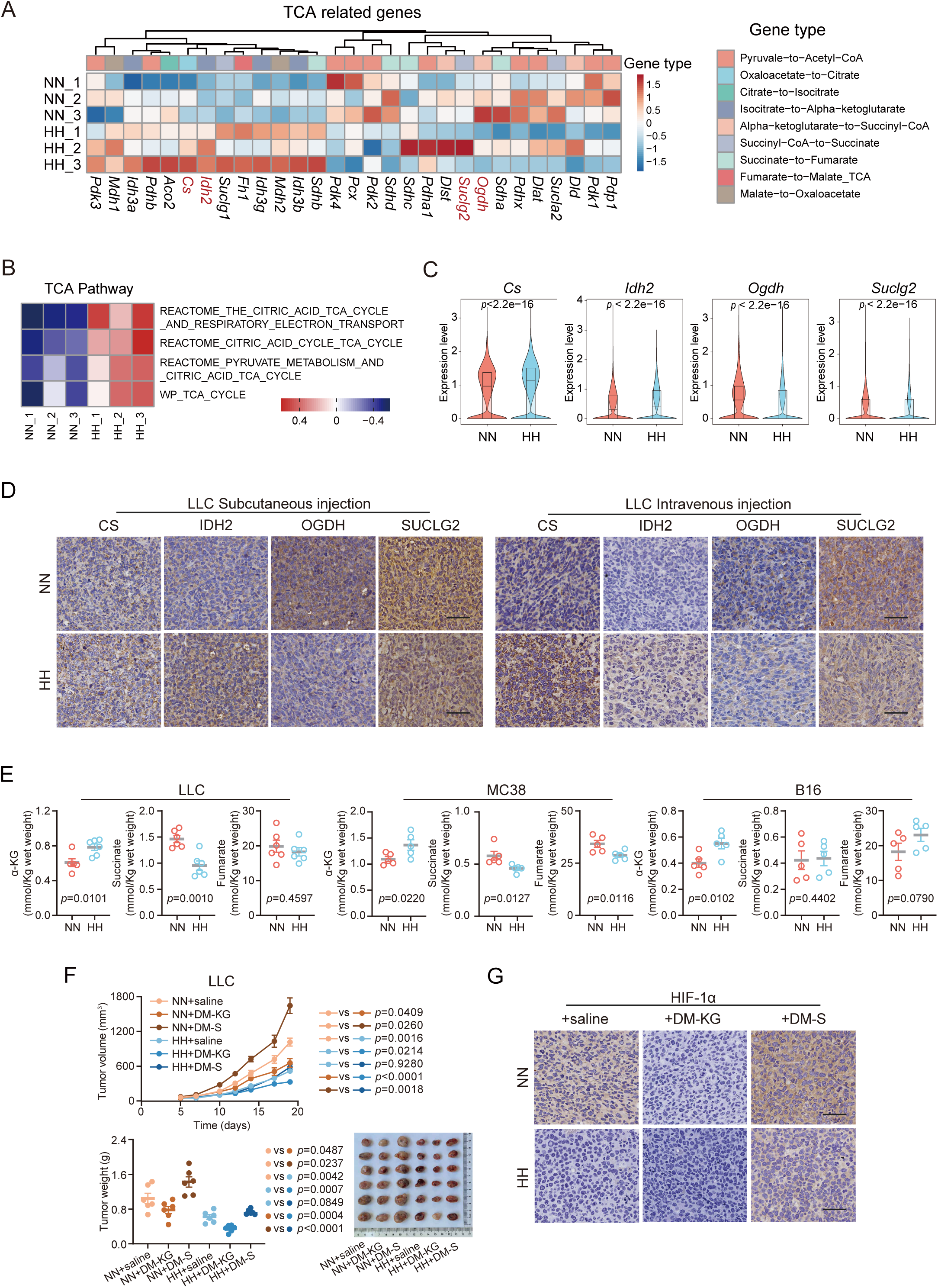
Reprogrammed TCA cycle promoted the HIF-1α pathway decline in tumor tissues under hypobaric hypoxia. (A) Gene expression profiles of TCA cycle-associated genes across all cells. (B) GSVA enrichment scores for TCA cycle-related pathways. (C) Violin plots depicting expression levels of TCA cycle enzymes *Cs*, *Idh2*, *Ogdh* and *Suclg2*. (D) Immunohistochemical staining of CS, IDH2, OGDH and SUCLG2 from LLC subcutaneous and intravenous tumor models. (E) Detection of α-KG, succinate and fumarate content in subcutaneous tumor tissues. (F) The growth curve, weight and images of tumor tissues from LLC subcutaneous tumor models receiving saline, DM-KG and DM-S treatment under plain and plateau. (G) Immunohistochemical staining of HIF-1α from LLC subcutaneous tumor models receiving saline, DM-KG and DM-S under plain and plateau. Data represent mean ± SEM. Scale bar=50 μm.

To prove the role of reprogrammed citrate cycle, α-KG and succinate were supplied into mice subcutaneously bearing LLC tumors. And succinate could reverse the inhibitory role and α-KG could enhance antitumor ability under hypobaric hypoxia. Moreover, the α-KG treated tumors under high altitude showed less volume and weight than that under plain, implying enhanced effect of hypobaric hypoxia on α-KG treatment. However, the tumors was promoted by succinate treatment (Figure 3F). More importantly, the level of HIF-1α was promoted by succinate treatment and compromised by α-KG treatment (Figure 3G). These results support that the antitumor effect of hypobaric hypoxia was dependent on remodeling citrate cycle induced HIF-1α inhibition.

### 3.4 Reprogramming citrate cycle of hypobaric hypoxia was transcriptionally regulated by TCF12 and YBX1

Given the results above, we could conclude that hypobaric hypoxia reshaped citrate cycle to regulate HIF-1α pathway. Since the mRNA and protein levels of Cs, Idh2, Ogdh and Suclg2 were affected, we supposed that these metabolic enzymes were transcriptionally regulated by high altitude exposure. Therefore, we investigated transcription factors with significant changes in tumor cells under hypoxic conditions. Our analysis revealed transcription factors with significantly enhanced activity (Figure 4A) and markedly weakened activity (Figure B). Furthermore, combination with JASPAR PWMs motif prediction of binding sites, *Ybx1* and *Tcf12* was recognized to regulate *Idh2* and *Ogdh* expression respectively. Moreover, TCF12 was decreased and YBX1 was increased in tumor tissues with hypobaric hypoxia treatment (Figure 4C). And overexpression *Ybx1* could promote *Idh2* expression (Figure 4D) and silencing *Tcf12* could downregulate the expression of *Ogdh* (Figure 4E). Furthermore, the results of luciferase reporter gene assays proved that YBX1 and TCF12 transcriptionally regulate the expression of *Idh2* and *Ogdh* respectively (Figure 4D-E). However, the mechanism of CS and SUCLG2 transformation by hypobaric hypoxia remains to be clarified.

**Figure 4.**
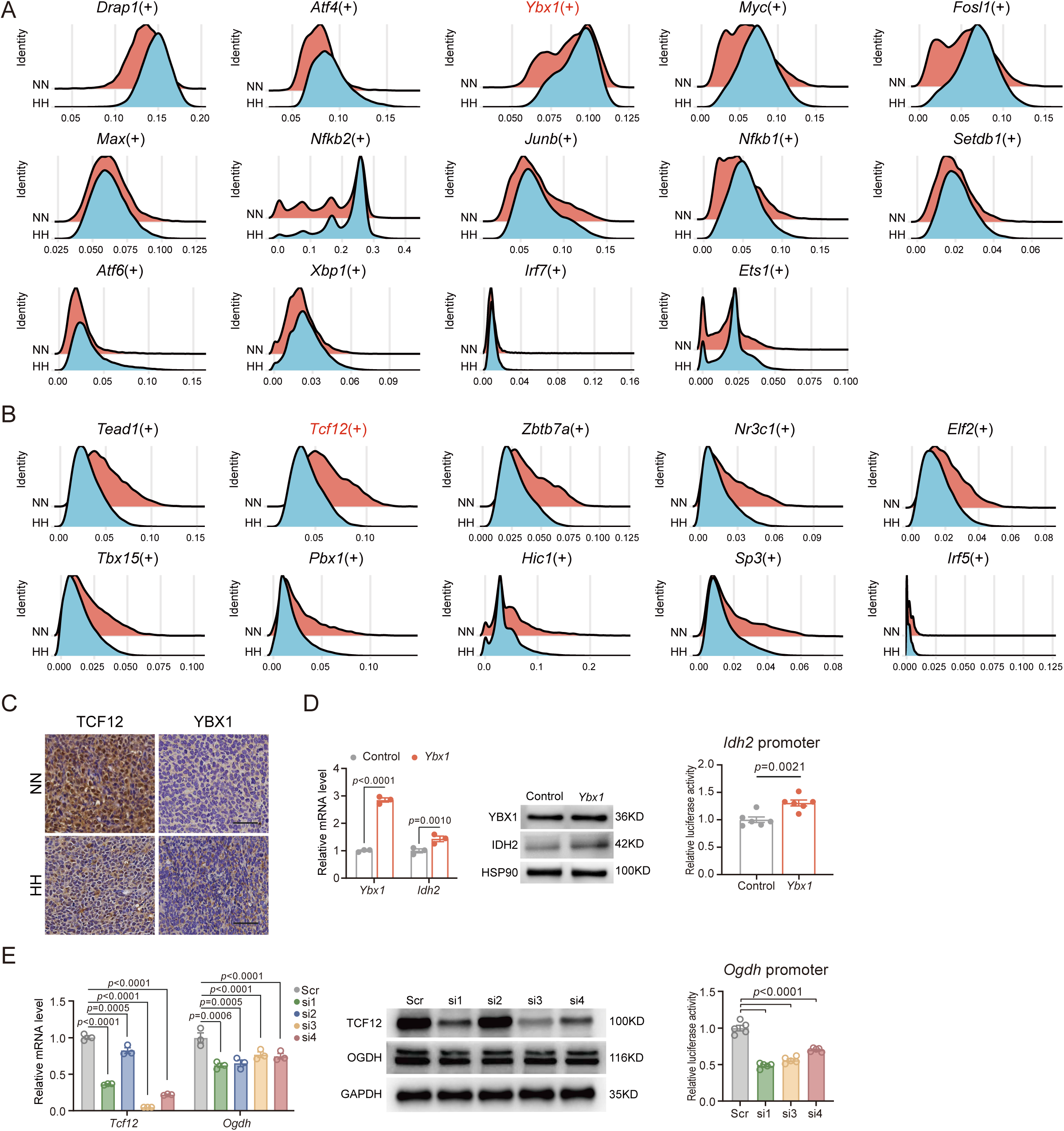
Reprogrammed TCA cycle under hypobaric hypoxia was transcriptionally regulated by YBX1 and TCF12. Transcriptional factors with significantly enhanced activity (A) and markedly weakened activity (B) observed under hypobaric hypoxia in tumor cells. (C) Immunohistochemical staining of YBX1 and TCF12 from LLC subcutaneous tumor models. The expression levels and promoter activity of *Idh2* with *Ybx1* vector transfection (D), and the expression levels and promoter activity of *Ogdh* with *Tcf12* siRNA transfection (E). Data represent mean ± SEM. Scale bar=50 μm.

### 3.5 Hypobaric hypoxia presented beneficial effect on tumor adjuvant therapy

To optimize the condition of hypobaric hypoxia treatment, we evaluated the effect of hypobaric hypoxia with different altitudes and periods on the tumor progression. And 5,800m high altitude exposure for 4h every day still showed significant inhibitory role on LLC subcutaneous bearing tumors (Figure 5A). Moreover, the impact of this treatment on the citrate cycle was consistent with that of 5,800m high altitude exposure for 24h every day (Figure 5B).

**Figure 5.**
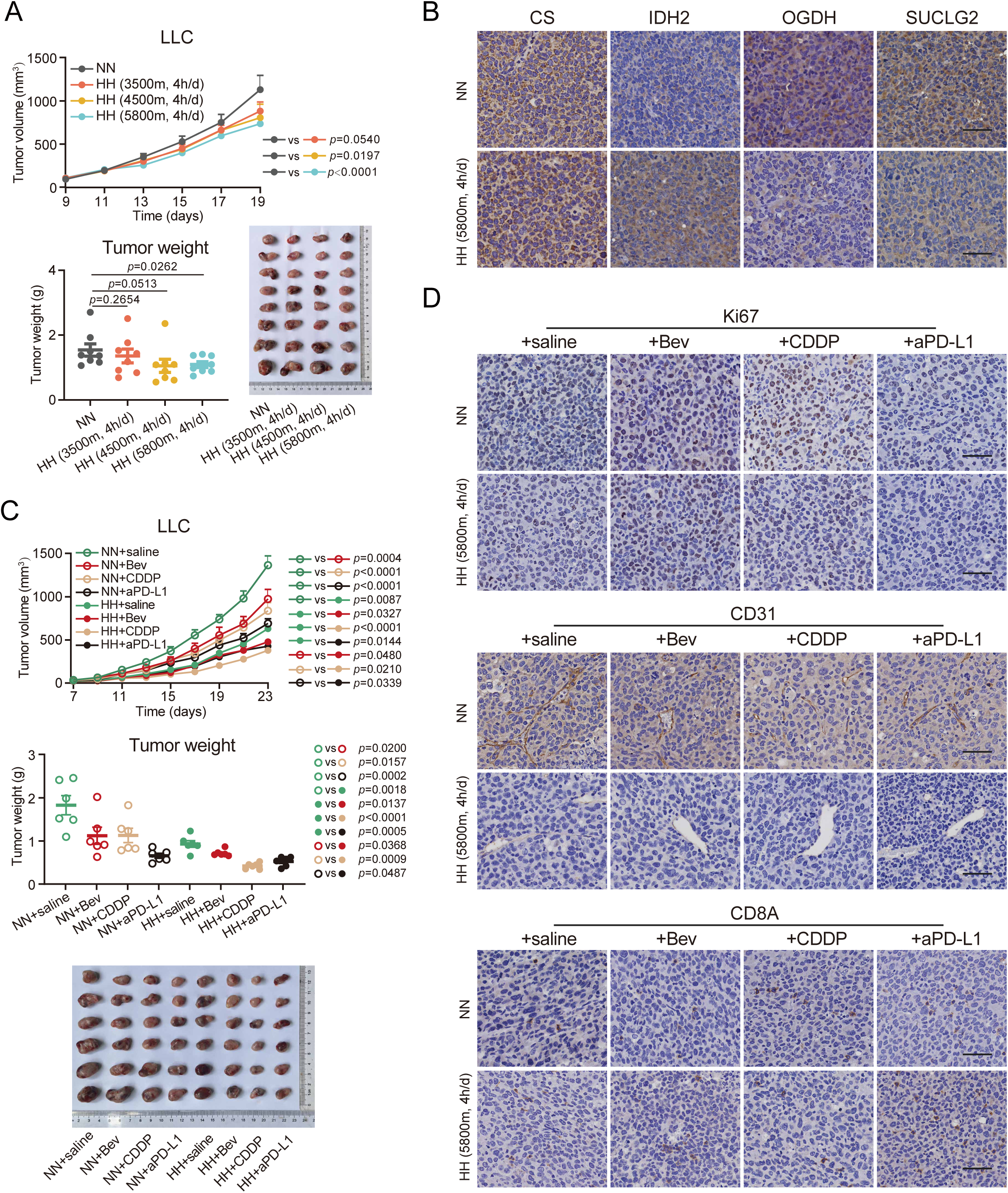
Hypobaric hypoxia improved tumor adjuvant therapies. (A) The growth curve, weight and images of tumor tissues from LLC subcutaneous tumor models under normobaric normoxia, 3,500m 4h/d, 4,500m 4h/d and 5,800m 4h/d. (B) Immunohistochemical staining of CS, IDH2, OGDH and SUCLG2 from LLC subcutaneous tumor models under plain and 5,800m 4h/d. (C) The growth curve, weight and images of tumor tissues from LLC subcutaneous tumor models receiving saline, bevacizumab (Bev), CDDP and anti-PD-L1 (αPD-L1) treatment under plain and 5,800m 4h/d. (D) Immunohistochemical staining of Ki67, CD31 and CD8A from LLC subcutaneous tumor models receiving saline, Bev, CDDP and αPD-L1 treatment under plain and 5,800m 4h/d. Data represent mean ± SEM. Scale bar=50 μm.

To investigate the effect of hypobaric hypoxia treatment on current clinical adjuvant therapies, we observed the tumor progression treated with bevacizumab, CDDP and anti PD-L1 (αPD-L1) under plain and plateau (5,800m 4h/d) condition. The results showed that hypobaric hypoxia could significantly enhance the antitumor ability of bevacizumab, CDDP and αPD-L1 (Figure 5C). Moreover, this treatment could display beneficial effect on these adjuvant therapies by downregulation of Ki67 and CD31 and upregulation of CD8A (Figure 5D). These results suggest that hypobaric hypoxia displays a great potential to be a new adjuvant therapy for tumor treatment.

## 4 Discussion

Hypoxia is a pivotal factor for tumor progression, triggering genetic, transcriptional, translational and epigenetic adaptations closely related to therapy resistance, metastasis and patient mortality [24]. Moreover, hypoxia creates immunosuppressive tumor microenvironment through reducing number and function of effector cells and production and release of effector cytokines, supporting immunosuppressive cells and increasing production and release of immunosuppressive cytokines [25–26]. And hypoxia sensing signaling plays critical roles in the modulation of cancer progression. The key molecules of the hypoxia sensing signaling include HIFs, which widely controls oxygen responsive genes, the central members of the 2-oxoglutarate (2-OG)-dependent dioxygenases, such as PHDs, and VHL. Among these molecules, HIFs are widely involved into the various oncogenic processes above [27]. Therefore, tumor vascular normalization or limitation of hypoxia induced pathways show a great potential for antitumor treatment [28]. Moreover, oxygen supplementation, such as hyperbaric oxygen and respiratory hyperoxia, showed antitumor effects by various pathways [8–9]. According to these research, rich oxygen supply systemically or locally into tumor displayed repressive role for tumor.

Whereas, environmental oxygen deprivation also presents antioncogenic ability. The incidence and mortality of many cancers in high altitude region lowers than that in plain area [5–6]. A research of calculating cancer incidence in Western United States showed lung cancer incidence decreased with altitude elevation and considered oxygen as an inhaled carcinogen [29]. And our work presented the inhibitory role of hypobaric hypoxia treatment for tumor progression. Different from basic research with hypoxia inducing HIF-1α expression, hypobaric hypoxia presented the suppressive role on HIF-1α pathway. However, hypobaric hypoxia doesn’t certainly represent local hypoxia in tumor tissues, which is induced by inadequate vascular formation. As to the clinical studies from high altitude, there are no definitive data that people in high altitude show high HIF-1α expression [30]. And a study showed the co-adapted Tibetan-specific haplotype encoding for PHD2 showed promotive effect on HIF-1α degradation under hypoxic conditions, which induced HIF-1α reduction [31]. In that case, Tibetan should show lower HIF-1α, consistently with decreased tumor incidence and mortality. However, a recent study considered that HIF-2α played a similar role in hypoxia adaptation between high altitude populations and tumors. However, this relationship was currently identified only in sympathetic tumors instead of malignant tumors and the tumorigenic effects showed hypoxia tissue-specific and time-sensitive. Therefore, we referred that this convergence between high altitude adaptation and tumor progression under hypoxia could be only limited into pheochromocytomas or paragangliomas, but not commonly existed [7]. Moreover, genetic fitted genes under high oxygen and low oxygen mostly showed no connection to HIF/ROS, implying the complexity about oxygen sensing and response [32].

Metabolic reprogramming is widely regarded as a remarkable hallmark of tumors for supplying high energetics and metabolites. And many metabolites have been identified to be involved in tumor promotion or inhibition [33]. Wherein, α-KG, succinate and fumarate, regarded as intermediate metabolites in citrate cycle, have been reported to be involved in HIF-1α regulation. HIF-1α is ubiquitylated by VHL and degraded by proteasome, which is dependent on the hydroxylation of two proline residues in HIF-1α. The hydroxylation reaction, mainly catalyzed by PHD2, is coupled to oxidative decarboxylation of α-KG to succinate and carbon dioxide [34–35]. And hypoxia confines HIF-1α hydroxylation and ubiquitin mediated proteasomal degradation and promotes the expression of HIF-1α regulated genes, such as glycolytic enzymes, angiogenetic and immune tolerance proteins. In addition, the transcriptional activity of HIF-1α is finely tuned by FIH-1, which mediates the hydroxylation of asparaginyl residue in HIF-1α and prevents recruitment of co-activators p300/CBP [36]. According to our results, the expression of PHD2, VHL, FIH-1 and p300/CBP did not show noticeable alteration. Although PHD2 is a well characterized oxygen-sensitive protein and its activity is dominated by tissue oxygen levels, the oxygen levels in tumor tissues between normoxia and hypoxia show no significant difference. Importantly, the accumulation succinate and fumarate, components in TCA cycle, could inhibit PHD2 activity by competing with α-KG [37–38]. However, hypobaric hypoxia induced α-KG increase and succinate decrease, which supported the antitumor effect of high altitude exposure. Recently, Cui et al. developed an antitumor agent to promote α-KG production and HIF-1α degradation by IDH1 activation, which could be achieved by hypobaric hypoxia [39].

Besides regulating PHD2 activity, α-ketoglutarate, succinate and fumarate are greatly involved into tumor initiation and progression. α-KG was recognized as an effector of p53-mediated tumor suppression and could promote tumor differentiation, induce apoptosis and antioxidative capacity and attenuate malignant progression [40–42]. Combination of BCAT1 inhibition and α-KG triggered metabolic synthetic lethality in glioblastoma [43]. Moreover, α-KG accumulation could overcome immune evasion and improve immunotherapy [44–45]. However, α-KG was also reported to promote immune escape by suppressing antigen presentation in cholangiocarcinoma [46] and accelerating M2 macrophage activation and proliferation in multiple cancers [47]. Different from α-KG, succinate was widely recognized as an oncometabolite. Succinate promoted cancer development through regulation of growth, epithelial to mesenchymal transition, cell migration invasion, angiogenesis, chemotherapy resistance and cancer cell stemness [48–51]. Moreover, succinate was involved in T cell exhaustion, immune-escape, immunotherapy resistance, macrophage polarization and cancer metastasis [52–54]. However, several studies reported the inhibitory effect of succinate on tumor progression through maintain the function of CD8^+^ T cells and promotion of pyroptosis, antigen presentation and classical M1-like polarization of macrophages [55–58]. Fumarate promoted tumor progression by PTEN inhibition and immune tolerance promotion [59–60]. Additionally, as competitive inhibitors, succinate and fumarate could inhibit α-ketoglutarate-dependent dioxygenases, which regulated histone modifications [61]. Then, the modulation of these metabolites could suppress tumor development through epigenetic regulation, which should be studied with further investigation. Generally speaking, besides with mediating HIF-1α downregulation, reprogramming of these metabolites could be involved in the antitumor activity through many pathways under hypobaric hypoxia. The production and degradation of α-KG and succinate is regulate by several critical metabolic enzymes in TCA cycle, which are greatly involved in various biological regulation [62–64]. In our work, metabolic reprogramming induced by hypobaric hypoxia was mainly regulated by CS, IDH2, OGDH and SUCLG2.

Hypobaric hypoxia treatment could enhance the efficacy of adjuvant therapy. And in the combined treated experiments, the drug doses within these adjuvant therapies was cut half and the efficacy of hypobaric hypoxia plain was close to that of other treatments under normoxic environment. Moreover, based on the gain effect to these therapies, hypobaric hypoxia displays a great potential for clinical antitumor therapy and will significantly decrease the therapeutic dose and side effect of these therapies above. However, the best optimal combined plans should be investigated and the mechanism for the enhanced effects will be clarified in the future.

This study demonstrated that hypobaric hypoxia presented the antitumor effect through suppressed malignant phenotype and improved immune response. Mechanically, HIF-1α pathway was significantly downregulated by α-KG accumulation and succinate reduction in the tumor environment, mediated by CS and IDH2 upregulation and OGDH and SUCLG2 downregulation in tumor cells under high altitude exposure (Figure 6). Furthermore, hypobaric hypoxia showed enhanced effect on antitumor adjuvant therapies and displayed a great value for tumor treatment.

**Figure 6.**
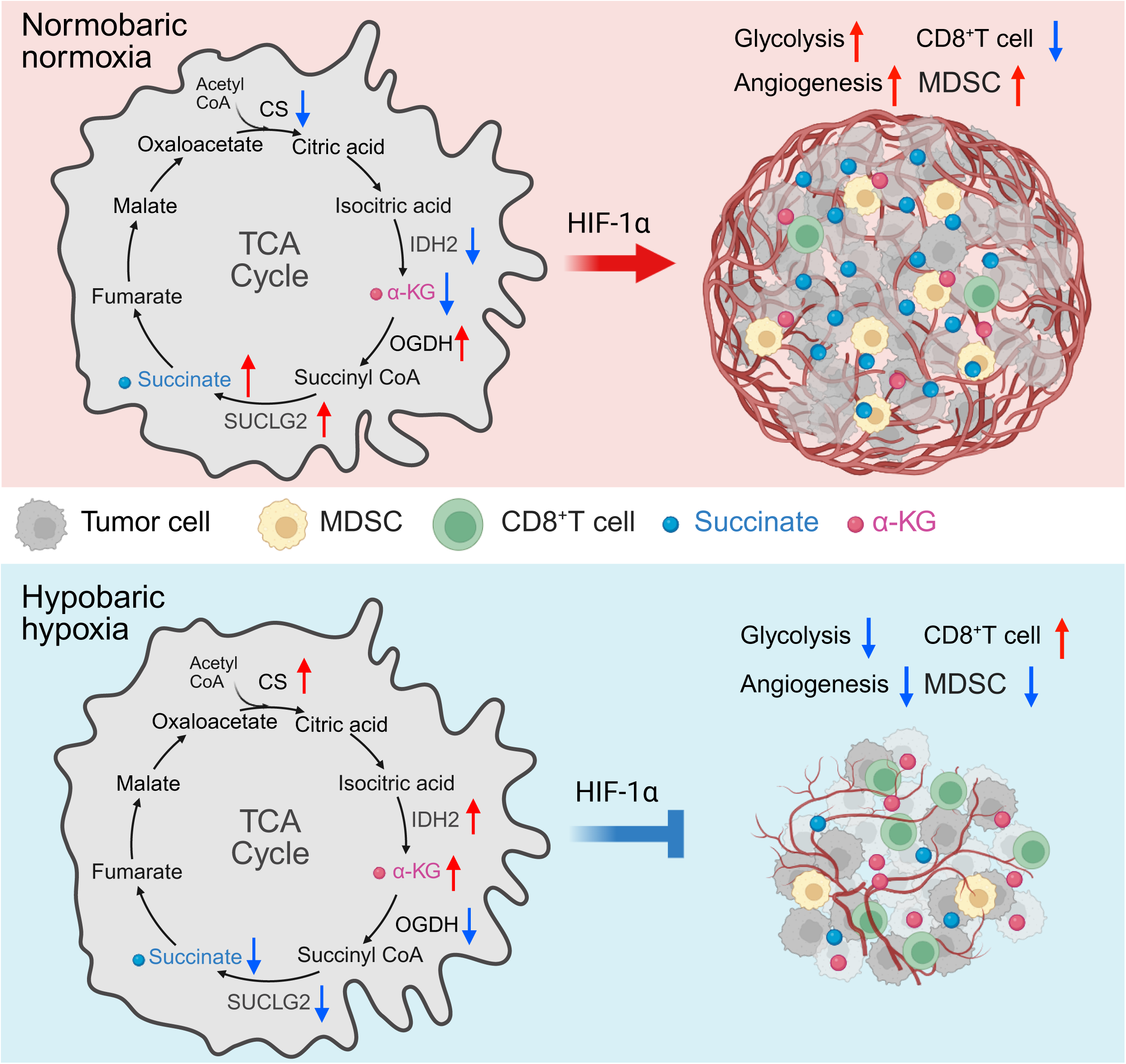
Schematic diagram of hypobaric hypoxia suppressing tumor progression. Tumor cells increase α-KG accumulation and decrease succinate release in tumor microenvironment mediated by CS and IDH2 upregulation and OGDH and SUCLG2 downregulation under hypobaric hypoxia. Reprogrammed α-KG and succinate distribution suppresses HIF-1α pathway to inhibit tumor progression and improve immune response. This image was created with BioRender.com.

## Author contributions

Yixing Gao and Xiuju He performed experiments, analyzed the data and wrote the original draft. Guoji E, Yan Wang, Shirui Huang, Dan Cai, Yumei Deng, Ping Gan, Rui Feng, Qian Liang, Ziyi Tang, Gang Xu and Jian Chen perform experiments. Hongyu Xue, Qiutong Wang, Huanhuan Liu, Guibo Li, Shuang Zeng and Min Xie analyzed the data. Yuqi Gao, Jiao Gong and Erlong Zhang conceived the study, supervised the project and revised the mauscript. All authors read and approved the final manuscript.

## Acknowledgements

This research was supported by grants from the National Natural Science Foundation of China (81802797) and the Program of Chongqing Talents.

## Data availability statement

The datasets used and/or analyzed during the current study are available from the corresponding author on reasonable request.

## Ethics statement

The animal experimental protocols were approved by the Animal Care and Use Committee Guidelines of the Army Medical University (AMUWEC20245324).

## Supplementary legends

**Figure S1.** (A) Images and weight of lung tissues from metastatic tumor models. Immunohistochemical staining of Ki67, CD31, CD8A and GZMB from MC38 and B16 subcutaneous (B) and MCA205 and B16 intravenous (C) tumor models. Data represent mean ± SEM. Scale bar=50 μm.

**Figure S2.** (A) Changes in cellular proportions from scRNA-seq. (B) Differentially expressed genes (DEGs) across distinct cell types. (C) *Hif-1*α and *Vegfa* expression profiles across cell types. (D) Expression patterns of HIF-1α downstream pathways associated with proliferation, invasion/metastasis, and cancer stem cells (CSCs). (E) Systematic subclassification of macrophage subsets. (F) FeaturePlot visualizing key marker genes for MDSC. (G) Comparative proportions of MDSC from scRNA-seq. \**p* < 0.05; \*\**p* < 0.01; \*\*\**p* < 0.001; \*\*\*\**p* < 0.0001.

**Figure S3.** (A) Heatmap depicting GSVA scores for hallmark gene sets under hypobaric hypoxia versus normobaric normoxia. (B) AddModuleScore analysis assessing Hypoxia, Peroxisome, Reactive oxygen species pathway, Inflammatory response, Apoptosis, and Il6-Jak-Stat3 signaling pathways across distinct cell types. \**p* < 0.05; \*\**p* < 0.01; \*\*\**p* < 0.001; \*\*\*\**p* < 0.0001.

**Figure S4.** Immunohistochemical staining of HIF-1α, HK1, VEGFA and CTLA4 from MC38 and B16 subcutaneous (A) and MCA205 and B16 intravenous (B) tumor models. (C) Immunohistochemical staining of Ki67, CD31 and CD8A from LLC subcutaneous tumor models receiving saline, DMOG and PX478 treatment under plain and plateau. Scale bar=50 μm.

**Figure S5.** (A) Levels of *Hif-1*α and *Hif-2*α mRNA in tumor tissues from subcutaneous models. (B) Immunohistochemical staining of VHL, FIH-1, PHD2, p300 and CA9 from LLC subcutaneous tumor models. (C) The content of ROS in tumor tissues from LLC subcutaneous tumor models. Immunohistochemical staining of CS, IDH2, OGDH and SUCLG2 from MC38 and B16 subcutaneous (D) and MCA205 and B16 intravenous (E) tumor models. Data represent mean ± SEM. Scale bar=50 μm.

**Figure S6.** (A) Immunohistochemical staining of ACO2, IDH3A, IDH3B, IDH3G, DLST, DLD, SUCLG1, SUCLA2, SDHA, SDHB, SDHC, SDHD, FH1, MDH1 and MDH2 from LLC subcutaneous tumor models. (B) Immunohistochemical staining of SDHA and FH1 from MC38 and B16 subcutaneous tumor models. Scale bar=50 μm.

**Table S1.** Comprehensive details of the sequencing methodologies and analytical procedures.

**Table S2.** Information of qPCR primers and RNA oligos used in this study.

**Table S3.** Information of primary antibodies used in this study.

## References

1. Kakkad S, Krishnamachary B, Jacob D, Pacheco-Torres J, Goggins E, Bharti SK, et al. Molecular and functional imaging insights into the role of hypoxia in cancer aggression. Cancer Metastasis Rev. 2019;38(1-2):51–64.

2. Chen Z, Han F, Du Y, Shi H, Zhou W. Hypoxic microenvironment in cancer: molecular mechanisms and therapeutic interventions. Signal Transduct Target Ther. 2023;8(1):70.

3. Rey-Keim S, Schito L. Origins and molecular effects of hypoxia in cancer. Semin Cancer Biol. 2024;106–107:166-178.

4. Abu El, Maaty MA, Terzic J, Keime C, Rovito D, Lutzing R, Yanushko D, et al. Hypoxia-mediated stabilization of HIF1A in prostatic intraepithelial neoplasia promotes cell plasticity and malignant progression. Sci Adv. 2022;8(29):eabo2295.

5. Thiersch M, Swenson ER. High altitude and cancer mortality. High Alt Med Biol. 2018;19(2):116–123.

6. Calderon-Gerstein WS, Torres-Samaniego G. High altitude and cancer: an old controversy. Respir Physiol Neurobiol. 2021;289:103655.

7. Arenillas C, Celada L, Ruiz-Cantador J, Calsina B, Datta D, Garcia-Galea E, et al. Convergent genetic adaptation in human rumors developed under systemic hypoxia and in populations living at high altitudes. Cancer Discov. 2025;15(5):1037–1062.

8. Hatfield SM, Kjaergaard J, Lukashev D, Schreiber TH, Belikoff B, Abbott R, et al. Immunological mechanisms of the antitumor effects of supplemental oxygenation. Sci Transl Med. 2015;7(277):277ra30.

9. Deng Q, Yang X, Li Z. Hyperbaric oxygen: a multifaceted approach in cancer therapy. Med Gas Res. 2024;14(3):130–132.

10. Machado VF, da Rocha JJR, Parra RS, Feitosa MR, Leite CA, Minto SB, et al. Hyperbaric oxygen therapy increases the effect of 5-fluorouracil chemotherapy on experimental colorectal cancer in mice. Med Gas Res. 2024;14(3):121–126.

11. Xiao C, Li J, Hua A, Wang X, Li S, Li Z, et al. Hyperbaric oxygen boosts antitumor efficacy of copper-diethyldithiocarbamate nanoparticles against pancreatic ductal adenocarcinoma by regulating cancer stem cell metabolism. Research (Wash D C). 2024;7:0335.

12. Xiong Y, Yong Z, Xu C, Deng Q, Wang Q, Li S, et al. Hyperbaric oxygen activates enzyme-driven cascade reactions for cooperative cancer therapy and cancer stem cells elimination. Adv Sci (Weinh). 2023;10(21):e2301278.

13. Yu L, Hales CA. Long-term exposure to hypoxia inhibits tumor progression of lung cancer in rats and mice. BMC Cancer. 2011;11:331.

14. Zhang S, Hu X, Sun M, Chen X, Le S, Wang X, et al. Potential role of hypobaric hypoxia environment in treating pan-cancer. Sci Rep. 2025;15(1):12942.

15. McGinnis CS, Murrow LM, Gartner ZJ. DoubletFinder: doublet detection in single-cell RNA sequencing data using artificial nearest neighbors. Cell Syst. 2019;8(4):329–337.e4.

16. Hao Y, Hao S, Andersen-Nissen E, Mauck WM 3rd, Zheng S, Butler A, et al. Integrated analysis of multimodal single-cell data. Cell. 2021;184(13):3573–3587.e29.

17. Korsunsky I, Millard N, Fan J, Slowikowski K, Zhang F, Wei K, et al. Fast, sensitive and accurate integration of single-cell data with Harmony. Nat Methods. 2019;16(12):1289–1296.

18. Yu G, Wang LG, Han Y, He QY. clusterProfiler: an R package for comparing biological themes among gene clusters. OMICS. 2012 May;16(5):284–287.

19. Xu S, Hu E, Cai Y, Xie Z, Luo X, Zhan L, et al. Using clusterProfiler to characterize multiomics data. Nat Protoc. 2024;19(11):3292–3320.

20. Hanzelmann S, Castelo R, Guinney J. GSVA: gene set variation analysis for microarray and RNA-seq data. BMC Bioinformatics. 2013;14:7.

21. Aibar S, Gonzalez-Blas CB, Moerman T, Huynh-Thu VA, Imrichova H, Hulselmans G, et al. SCENIC: single-cell regulatory network inference and clustering. Nat Methods. 2017;14(11):1083–1086.

22. Van de Sande B, Flerin C, Davie K, De Waegeneer M, Hulselmans G, Aibar S, et al. A scalable SCENIC workflow for single-cell gene regulatory network analysis. Nat Protoc. 2020;15(7):2247–2276.

23. Choudhry H, Harris AL. Advances in hypoxia-inducible factor biology. Cell Metab. 2018;27(2):281–298.

24. Lee P, Chandel NS, Simon MC. Cellular adaptation to hypoxia through hypoxia inducible factors and beyond. Nat Rev Mol Cell Biol. 2020;21(5):268–283.

25. Fan P, Zhang N, Candi E, Agostini M, Piacentini M, TOR Centre et al. Alleviating hypoxia to improve cancer immunotherapy. Oncogene. 2023;42(49):3591–3604.

26. Jing X, Yang F, Shao C, Wei K, Xie M, Shen H, et al. Role of hypoxia in cancer therapy by regulating the tumor microenvironment. Mol Cancer. 2019;18(1):157.

27. Yang G, Shi R, Zhang Q. Hypoxia and oxygen-sensing signaling in gene regulation and cancer progression. Int J Mol Sci. 2020;21(21):8162.

28. Zheng R, Li F, Li F, Gong A. Targeting tumor vascularization: promising strategies for vascular normalization. J Cancer Res Clin Oncol. 2021;147(9):2489–2505.

29. Simeonov KP, Himmelstein DS. Lung cancer incidence decreases with elevation: evidence for oxygen as an inhaled carcinogen. PeerJ. 2015;3:e705.

30. Arciero E, Almarri MA, Mezzavilla M, Xue Y, Hallast P, Hammoud C, et al. Whole-genome sequences provide insights into the formation and adaptation of human populations in the Himalayas. Curr Biol. 2025:S0960-9822(25)00808-5.

31. Lanikova L, Reading NS, Hu H, Tashi T, Burjanivova T, Shestakova A, et al. Evolutionary selected Tibetan variants of HIF pathway and risk of lung cancer. Oncotarget. 2017;8(7):11739–11747.

32. Jain IH, Calvo SE, Markhard AL, Skinner OS, To TL, Ast T, et al. Genetic screen for cell fitness in high or low oxygen highlights mitochondrial and lipid metabolism. Cell. 2020;181(3):716–727.e11.

33. Pavlova NN, Zhu J, Thompson CB. The hallmarks of cancer metabolism: Still emerging. Cell Metab. 2022;34:355–77.

34. Semenza GL. Hypoxia-inducible factors in physiology and medicine. Cell. 2012;148(3):399–408.

35. Kaelin WG Jr, Ratcliffe PJ. Oxygen sensing by metazoans: the central role of the HIF hydroxylase pathway. Mol Cell. 2008;30(4):393–402.

36. Lando D, Peet DJ, Whelan DA, Gorman JJ, Whitelaw ML. Asparagine hydroxylation of the HIF transactivation domain a hypoxic switch. Science. 2002;295(5556):858–861.

37. Pollard PJ, Briere JJ, Alam NA, Barwell J, Barclay E, Wortham NC, et al. Accumulation of Krebs cycle intermediates and over-expression of HIF1alpha in tumours which result from germline FH and SDH mutations. Hum Mol Genet. 2005;14(15):2231–2239.

38. Xu W, Yang H, Liu Y, Yang Y, Wang P, Kim SH, et al. Oncometabolite 2-hydroxyglutarate is a competitive inhibitor of alpha-ketoglutarate-dependent dioxygenases. Cancer Cell. 2011;19(1):17–30.

39. Cui Z, Li C, Liu W, Sun M, Deng S, Cao J, et al. Scutellarin activates IDH1 to exert antitumor effects in hepatocellular carcinoma progression. Cell Death Dis. 2024;15(4):267.

40. Morris JP 4th, Yashinskie JJ, Koche R, Chandwani R, Tian S, Chen CC, et al. alpha-Ketoglutarate links p53 to cell fate during tumour suppression. Nature. 2019;573(7775):595–599.

41. Tran TQ, Hanse EA, Habowski AN, Li H, Ishak Gabra MB, Yang Y, et al. alpha-Ketoglutarate attenuates Wnt signaling and drives differentiation in colorectal cancer. Nat Cancer. 2020;1(3):345–358.

42. Greilberger J, Herwig R, Kacar M, Brajshori N, Feigl G, Stiegler P, et al. Alpha-ketoglutarate or 5-hmf: single compounds effectively eliminate leukemia cells via caspase-3 apoptosis and antioxidative pathways. Int J Mol Sci. 2022;23(16):9034.

43. Zhang B, Peng H, Zhou M, Bao L, Wang C, Cai F, et al. Targeting BCAT1 combined with alpha-ketoglutarate triggers metabolic synthetic lethality in glioblastoma. Cancer Res. 2022;82(13):2388–2402.

44. Li L, Zeng X, Chao Z, Luo J, Guan W, Zhang Q, et al. Targeting alpha-ketoglutarate disruption overcomes immunoevasion and improves pd-1 blockade immunotherapy in renal cell carcinoma. Adv Sci (Weinh). 2023;10(27):e2301975.

45. Liang L, Kuang X, He Y, Zhu L, Lau P, Li X, et al. Alterations in PD-L1 succinylation shape anti-tumor immune responses in melanoma. Nat Genet. 2025;57(3):680–693.

46. Zhang N, Sun L, Zhou S, Ji C, Cui T, Chu Q, et al. Cholangiocarcinoma PDHA1 succinylation suppresses macrophage antigen presentation via alpha-ketoglutaric acid accumulation. Nat Commun. 2025;16(1):3177.

47. Cai Z, Li W, Hager S, Wilson JL, Afjehi-Sadat L, Heiss EH, et al. Targeting PHGDH reverses the immunosuppressive phenotype of tumor-associated macrophages through alpha-ketoglutarate and mTORC1 signaling. Cell Mol Immunol. 2024;21(5):448–465.

48. Tong Y, Qi Y, Xiong G, Li J, Scott TL, Chen J, et al. The PLOD2/succinate axis regulates the epithelial-mesenchymal plasticity and cancer cell stemness. Proc Natl Acad Sci U S A. 2023;120(20):e2214942120.

49. Dalla Pozza E, Dando I, Pacchiana R, Liboi E, Scupoli MT, Donadelli M, et al. Regulation of succinate dehydrogenase and role of succinate in cancer. Semin Cell Dev Biol. 2020;98:4–14.

50. Kuo CC, Wu JY, Wu KK. Cancer-derived extracellular succinate: a driver of cancer metastasis. J Biomed Sci. 2022;29(1):93.

51. Yuan Y, Wu Y, Li C, Huang Z, Peng D, Wu Z, et al. Circ0515 reprogramming mitochondrial succinate metabolism and promotes lung adenocarcinoma progression through regulating SDHB. Cell Death Dis. 2025;16(1):497.

52. Jiang SS, Xie YL, Xiao XY, Kang ZR, Lin XL, Zhang L, et al. Fusobacterium nucleatum-derived succinic acid induces tumor resistance to immunotherapy in colorectal cancer. Cell Host Microbe. 2023;31(5):781–797.e9.

53. Wu JY, Huang TW, Hsieh YT, Wang YF, Yen CC, Lee GL, et al. Cancer-derived succinate promotes macrophage polarization and cancer metastasis via succinate receptor. Mol Cell. 2020;77(2):213–227.e5.

54. Ma S, Ong LT, Jiang Z, Lee WC, Lee PL, Yusuf M, et al. Targeting P4HA1 promotes CD8(+) T cell progenitor expansion toward immune memory and systemic anti-tumor immunity. Cancer Cell. 2025;43(2):213–231.e9.

55. Zheng P, Wang G, Liu B, Ding H, Ding B, Lin J. Succinate nanomaterials boost tumor immunotherapy via activating cell pyroptosis and enhancing MHC-I expression. J Am Chem Soc. 2025;147(2):1508–1517.

56. Mangalhara KC, Varanasi SK, Johnson MA, Burns MJ, Rojas GR, Esparza Molto PB, et al. Manipulating mitochondrial electron flow enhances tumor immunogenicity. Science. 2023;381(6664):1316–1323.

57. Elia I, Rowe JH, Johnson S, Joshi S, Notarangelo G, Kurmi K, et al. Tumor cells dictate anti-tumor immune responses by altering pyruvate utilization and succinate signaling in CD8(+) T cells. Cell Metab. 2022;34(8):1137–1150.e6.

58. Lu S, Li J, Li Y, Liu S, Liu Y, Liang Y, et al. Succinate-loaded tumor cell-derived microparticles reprogram tumor-associated macrophage metabolism. Sci Transl Med. 2025;17(793):eadr4458.

59. Ge X, Li M, Yin J, Shi Z, Fu Y, Zhao N, et al. Fumarate inhibits PTEN to promote tumorigenesis and therapeutic resistance of type2 papillary renal cell carcinoma. Mol Cell. 2022;82(7):1249–1260.e7.

60. Cheng J, Yan J, Liu Y, Shi J, Wang H, Zhou H, et al. Cancer-cell-derived fumarate suppresses the anti-tumor capacity of CD8(+) T cells in the tumor microenvironment. Cell Metab. 2023;35(6):961–978.e10.

61. Brunner JS, Finley LWS. Metabolic determinants of tumour initiation. Nat Rev Endocrinol. 2023;19(3):134–150.

62. Chaves-Perez A, Millman SE, Janaki-Raman S, Ho YJ, Hinterleitner C, Barthet VJA, et al. Metabolic adaptations direct cell fate during tissue regeneration. Nature. 2025;643(8071):468–477.

63. Jaccard A, Wyss T, Maldonado-Perez N, Rath JA, Bevilacqua A, Peng JJ, et al. Reductive carboxylation epigenetically instructs T cell differentiation. Nature. 2023;621(7980):849–856.

64. Wu MJ, Kondo H, Kammula AV, Shi L, Xiao Y, Dhiab S, et al. Mutant IDH1 inhibition induces dsDNA sensing to activate tumor immunity. Science. 2024;385(6705):eadl6173.

